# Ketone ester supplementation reduces age-associated B cell activation in aged mice without impairing antibody responses

**DOI:** 10.64898/2026.04.20.718782

**Authors:** Ariel Adkisson-Floro, Ritesh Tiwari, Mitsunori Nomura, Rebeccah R. Riley, Ryan Kwok, Durai Sellegounder, Mir M. Khalid, Herbert G. Kasler, John C. Newman, Eric Verdin

## Abstract

**Background:** Aging in the immune system results in increased susceptibility to infections, exacerbated autoimmunity, and reduced responsiveness to vaccines. However, there are no current established interventions for immune aging. Ketogenic diets and fasting have been researched as interventions against other aspects of aging and age-related diseases, and they work in part by increasing circulating levels of ketone bodies, which have anti-inflammatory properties and can boost T cell function. Exogenous ketones, such as ketone esters, are currently being studied as a more accessible approach to obtain the benefits of ketone bodies through direct supplementation. Here, we investigated whether ketone ester supplementation improves immune function during aging. Aged (19-month-old) C57BL/6JN mice were given a diet supplemented with the ketone ester or a control diet for 15 weeks.

**Results:** We found that the ketone ester diet decreased activation of B cells, especially age-associated B cells, in the spleen. The diet also reduced the proportion of age-associated B cells defined by CD11b and CD11c expression, though not of the broader CD21^−^CD23^−^ population. Despite this decrease in activation, there was no impairment in antibody production, as mice on the ketone ester diet mounted a normal response to nitrophenyl-ovalbumin immunization. The ketone ester diet also reduced glucose dependence and translation in subsets of age-associated B cells. Treating splenocytes from aged mice with β-hydroxybutyrate was sufficient to suppress B cell translation and reduced mTORC1 signaling, indicating that ketone bodies act directly on B cells. β-hydroxybutyrate also decreased expression of T-bet, the signature transcription factor driving the autoimmune functions of age-associated B cells.

**Conclusions:** Our study elucidates the effect of ketone esters on B cells in the context of aging and unveils a new immunoregulatory role of ketone bodies on B cells.

## Background

Aging in the immune system increases vulnerability to illness. These age-related immune changes include a decrease in naive T and B cells, chronic inflammation, and an increase in exhausted cells (1,2). As a result, older adults become more susceptible to infections, respond less effectively to vaccinations, and are at increased risk for cancer and autoimmune diseases (2–4). This increase in autoimmune disease is partly attributed to an increase in a subset of B cells known as “age-associated B cells” (ABCs). First identified in aged mice by their low expression of CD21 and CD23 or their expression of CD11b and CD11c (5,6), ABCs are elevated in autoimmune diseases, such as systemic lupus erythematosus (7,8). ABCs contribute to autoimmunity by secretion of proinflammatory cytokines, production of autoantibodies, and presentation of autoantigens to T cells(7–9). In young mice (3-4 months old), age-associated B cells are only a minor proportion of splenic B cells (about 5%), which then grows about 5-fold by 22 months of age (5,10).

While many interventions are being studied to ameliorate aspects of immune aging, none are currently established. Some of these interesting interventions are fasting and ketogenic diets, which are being studied to treat non-immunologic aspects of aging and age-related diseases, such as type 2 diabetes and Alzheimer’s disease (11–13). These diets are also being studied in the context of immune dysfunction, including autoimmune diseases; for example, both ketogenic diets and fasting-mimicking diets ameliorate experimental autoimmune encephalomyelitis in mice, a model of multiple sclerosis (14–16). One mechanism by which fasting and ketogenic diets achieve their beneficial effects is via the increase in circulating ketone bodies, which can be used as an alternative energy source during periods of hypoglycemia (17). Ketone bodies also affect immune cells. For example, β-hydroxybutyrate (BHB), the primary ketone body produced, inhibited the NLRP3 inflammasome in murine bone marrow-derived macrophages and human monocytes (18). BHB promoted M2 macrophage polarization in a mouse model of colitis, demonstrating anti-inflammatory properties, yet has also been shown to promote T cell function (19–21).

While the effects of ketone bodies on innate immune cells and T cells have been studied, their effects on B cells remain largely unexplored. Due to difficulty with adherence to fasting or ketogenic diets, particularly for older adults, exogenous ketone supplementation is currently being studied as a more accessible alternative to reap the benefits of ketone bodies (22,23). Exogenous ketone supplements include both ketone esters and ketone salts, though ketone esters are more widely favored because they cause a more robust increase in circulating ketone body levels and are associated with fewer adverse events, such as gastrointestinal side effects (24). Once ingested, ketone esters are broken down by esterases in the gut and converted into ketone bodies, providing a stable way to supplement them directly (25). However, it is unclear how ketone esters impact the immune system, especially in the context of aging.

In this study, we sought to understand the effect of ketone esters on the immune system in aged mice. We found that ketone ester supplementation through diet decreased activation of B cells, including age-associated B cells, in the spleen, yet it did not impair B cell function as shown by functional antibody production to a nitrophenyl-ovalbumin (NP-OVA) immunization. These results were accompanied by an increase in ketone bodies, and BHB alone was sufficient to inhibit translation within B cells. This work uncovers a new role for ketone bodies as metabolic regulators of B cell activation, acting more selectively than broad immunosuppression by leaving antibody production intact. Considering the role of age-associated B cells in autoimmunity, our findings raise the question of whether ketone esters could be an intervention for age-related autoimmune disease.

## Methods

### Mice

All mice were maintained in accordance with the guidelines set forth by the National Institutes of Health, and all experimental protocols were approved by the Buck Institute Institutional Animal Care and Use Committee (IACUC). The Buck Institute is accredited by the Association for Assessment and Accreditation of Laboratory Animal Care (AAALAC). The mice were maintained in a specific pathogen-free barrier facility under a 6:00 am to 6:00 pm light cycle. The facility maintains a temperature of 68-72°F and an ambient humidity level exceeding 30%. All mice were maintained in groups of up to five mice per cage. The cages were furnished with wooden bedding, a nestlet, and a house, and were changed every two weeks. The diet was provided in a recess of the metal wire lid situated at the upper portion of the cage and changed every two weeks. The wire lid was changed with the diet and water provided in a bottle. Male and female C57BL/6JN mice were obtained from the National Institute on Aging’s Aged Rodent Colonies at 19 months of age. Mice were given access to water and their appropriate diet ad libitum. One week after the mice acclimated to vivarium environmental conditions, they were all switched to a control diet (Teklad, TD.150345). One week later, half of the mice were then switched to a formulated ketone ester diet (Teklad, TD.240740), which contained 20% of the ketone ester, bis-octanoyl (R)-1,3-butanediol (26–28). Per-calorie macronutrient content for the diets is as follows: control diet (TD.150345), 77.7% carbohydrate, 12.6% fat, 9.7% protein; ketone ester diet (TD.240740), 51.1% carbohydrate, 39.2% fat, 9.7% protein. Weights of the mice were measured weekly. The study began with 29 mice (14 females, 15 males). 24 mice (10 females, 14 males) survived the duration of the immunization experiment. Tissues were collected at ZT2–ZT3 over two days from 23 mice as one mouse died between the end of the immunization experiment and tissue collection. Mice were euthanized with CO_2_ followed by cervical dislocation.

### Tissue processing

Tissues collected from mice include spleens, femurs, and plasma. Plasma was collected via cardiac puncture. Spleens were collected from mice and processed into single-cell suspension with phosphate-buffered saline (PBS) and a 70µm filter. Femurs were also collected from mice and cut at each end to then flush out the bone marrow using PBS. Red blood cells were lysed with 1x ammonium-chloride-potassium (ACK) buffer for 1-2 minutes and PBS was added up to 45 mL. Cells were spun down, supernatant was removed, and the cell pellet was resuspended in PBS for cell counting via hemocytometer. Spleens were classified as enlarged if gross splenomegaly was noted at dissection. Mice with enlarged spleens were removed from the spleen immunophenotyping and spleen SCENITH analyses. Of the 23 mice from which tissues were collected, 2 control and 5 ketone ester mice were excluded, leaving n = 8 per group for immunophenotyping. The proportion of mice with enlarged spleens was not significantly different between diet groups (Fisher’s exact test, p = 0.405).

### NP-OVA immunization

For primary immunization, nitrophenyl-ovalbumin (Biosearch Technologies #N-5051) was mixed in a 1:1 ratio in Freund’s incomplete adjuvant (ThermoFisher #77145) at 500µg/mL using a mechanical syringe. Each mouse was immunized with 50µg of NP-OVA intra-peritoneally. For secondary challenge, NP-OVA was resuspended in PBS (Corning #21-031-CV), and each mouse was injected with 50µg intra-peritoneally. Blood was collected retro-orbitally into heparin tubes at the indicated timepoints at ZT3 – ZT6. Tubes were spun down for 5 minutes at 2,500 x *g* at room temperature. The top clear layer of plasma was collected into new tubes and stored at −80°C for later analyses.

### Flow cytometry

Cells were transferred to 96-well V-bottom plates in PBS and stained with Live/Dead Blue (Fisher Scientific #L23105) for 10 minutes at 4°C. Cells were then incubated with surface antibodies for 30 minutes, spun down, and washed with PBS. Cells were fixed at room temperature for 10 minutes using the eBioscience Foxp3/Transcription Factor kit (ThermoFisher #00-5523-00). After washing with permeabilization buffer, cells were incubated with intracellular antibodies for 30 minutes. Cells were washed again, resuspended in PBS, and acquired on a Cytek Aurora flow cytometer. Data were analyzed using SpectroFlo v3.3.0 and FlowJo v10.10.0. To calculate absolute cell counts, the total cell count was first multiplied by the percentage of the “cells” gate from flow to give its absolute count. Each subsequent subset was then calculated by multiplying its percentage of parent population by the absolute count of its parent gate. For immunophenotyping, the antibodies used are listed in Tables S2-S3. For the SCENITH experiment on splenocytes from mice on control or ketone ester diet, a subset of the same antibodies from Table S2 was used. For the cell culture stimulation experiment with BHB treatment, a subset of the antibodies from Table S2 was used with the following swaps or additions: CD19 BV605 (BD Biosciences Clone 1D3 #563148), CD23 PE-Dazzle594 (BioLegend Clone B3B4 #101634), p-S6 PE (BD Biosciences Clone N7-548 #560433). For the age-associated B cell differentiation experiment, a subset of the antibodies from Table S2 was used with the following swaps or additions: CD23 PE-Dazzle594 (listed above), B220 FITC (BioLegend Clone RA3-6B2 #103206), and T-bet BV711 (BioLegend Clone 4B10 #644820).

### SCENITH

SCENITH was performed on splenocytes from a subset of the same aged mice used for immunophenotyping. Because tissues were collected over two days, mice designated for SCENITH were assigned to the second collection day in advance, giving 6 control and 6 ketone ester mice. After exclusion of mice with enlarged spleens (see Tissue processing), SCENITH analyses included n = 6 control and n = 4 ketone ester mice. The SCENITH kit was used according to manufacturer’s instructions (Gammaomics, Alexa Fluor 488). Each sample was split into five different wells in a 96-well V-bottom plates and 5 µL of each inhibitor (2-deoxyglucose, oligomycin) was added to the appropriate well. Samples incubated at 37°C for 10 minutes. 5 µL of puromycin was added to all wells and samples incubated at 37°C for 1 hour. Samples then proceeded through flow cytometry staining as described earlier with puromycin staining at the intracellular staining step.

### Cell culture

Splenocytes were isolated from mice and cultured in complete media (RPMI 1640 with L-glutamine, 10% fetal bovine serum, 1% penicillin-streptomycin, 1% sodium pyruvate). For the stimulation experiments, splenocytes were stimulated with LPS (Sigma #L2630) at 10 µg/mL for four days or overnight along with 5mM of R-BHB sodium salt (Sigma #298360-1G). For the age-associated B cell differentiation experiment, B cells were magnetically isolated from splenocytes using the EasySep Mouse Pan-B Cell Isolation Kit (STEMCELL #19844A) according to manufacturer’s instructions. 2×10^5^ B cells were plated per well of a 24-well plate in a total volume of 1 mL in age-associated B cell differentiation media for 3 days. Age-associated B cell differentiation media consisted of 500 ng/mL CL097 (InvivoGen #tlrl-c97), 50 ng/mL IL-21 (BioLegend #574502), 1 ng/mL IFNγ (BioLegend #575302), 1 µg/mL goat anti-mouse IgM F(ab’)_2_ (Jackson ImmunoResearch # 115-005-020), and 1 µg/mL anti-mouse CD40 (Southern Biotech #164501). Cells were then stained with the flow cytometry and SCENITH protocols as described earlier.

### ELISA

To measure antibodies against nitrophenyl (NP), high binding plates (Greiner #655061) were coated overnight at 4°C with NP-9-BSA (Biosearch #N-5050L) or NP-32-BSA (Biosearch #N-5050H) in coating buffer (NaHCO_3_, Na_2_CO_3_, pH 9.2) at 10 µg/mL. Plates were then blocked overnight at 4°C with 1% Blotto (Bio-Rad #1706404) in washing buffer. After washing with wash buffer, plasma samples from mice were added at the highest dilution of 1:1000 in dilution buffer (0.25% Blotto in wash buffer) and diluted down threefold to the lowest dilution of 1:30,000. Plates were incubated overnight at 4°C. After washing, IgG-HRP (Southern Biotech #1031-05) or IgM-HRP (Southern Biotech #1021-05) was added at 1:3000 in dilution buffer for 1 hour at room temperature and plates were washed. Plates were developed with substrate solution (0.5 mg/mL o-phenylenediamine, 9.8 mM H_2_O_2_ in citrate phosphate buffer) and reaction was stopped with H_2_SO_4_. Absorbances were measured with an Agilent BioTek Epoch Microplate Spectrophotometer. Absorbance at 630 nanometers was subtracted from absorbance at 492 nanometers.

### BHB measurements

Mice had ad libitum access to their assigned diet throughout the study and were not fasted prior to any blood collection. Thus, plasma for R-BHB measurement was collected in the ad libitum-fed state. Serial collections were performed within the same window of the light phase (ZT3-ZT6) to limit variation from circadian and feeding rhythms, and the final sample was collected by cardiac puncture at terminal tissue collection (ZT2-ZT3). R-BHB levels were measured from undiluted mouse plasma according to manufacturer’s instructions (Stanbio #2440058).

### Statistical analysis

Wilcoxon Mann-Whitney U tests were used to compare groups unless otherwise mentioned when conditions were appropriately met to use a T-test. These comparisons were not adjusted for multiple comparisons, and the spleen and bone marrow immunophenotyping analyses (Figures 1, S2-S4) were exploratory. For the ex vivo β-hydroxybutyrate treatment experiments, splenocytes from each mouse were used in both untreated and treated conditions, so paired T-tests were used. One-sided tests were used for the translation measurements in Figure 4C because the a priori hypothesis based on Figure 3 was that BHB would decrease translation. All other comparisons in these experiments were two-sided, as the pS6 analyses were exploratory and the direction of any effect was not predicted. Graphs were generated using *ggplot2* version 3.5.2 in R version 4.4.2 (29). For spleen immunophenotyping and for the SCENITH experiment on splenocytes from these mice, mice with enlarged spleens were removed from analysis. Splenomegaly (Fisher’s exact test, p = 0.405) was not different between diet groups, nor by sex (Fisher’s exact test, p > 0.999 for both sexes). Survival (Fisher’s exact test, p = 0.390) was not different between diet groups, nor by sex (Fisher’s exact test, p = 0.560 for females, p > 0.999 for males). For NP ELISAs, exclusion of mice that did not mount a primary response was specified in advance of the analysis. These non-responders were identified from titers at day 14 and removed from all subsequent timepoints. Error bars throughout represent standard error of the mean (s.e.m.).

**Figure 1:**
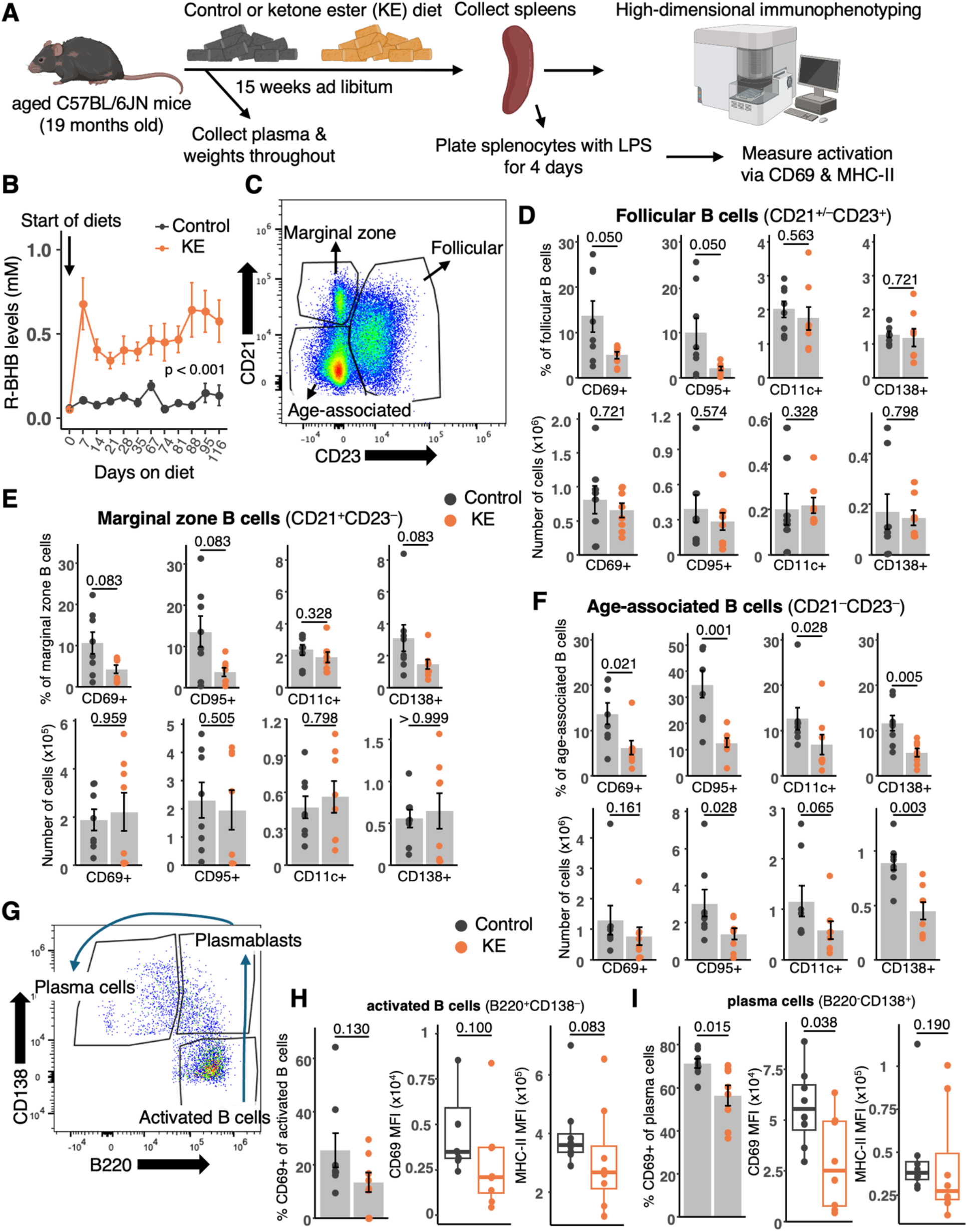
The ketone ester diet decreases activation of B cells. **A**) Experimental diagram for immunophenotyping on spleens from mice fed with the ketone ester (KE) diet or control diet and stimulating splenocytes with LPS (10 µg/mL) for 4 days to determine activation. Plasma was collected and weights were measured throughout the experiment. **B**) R-BHB levels in plasma from mice throughout the experiment (mean ± s.e.m.). A linear mixed effects model on log-transformed R-BHB levels, excluding baseline (day 0), was used to compare mice on control diet to mice on KE diet (p < 0.001). Data are plotted on original scale. **C**) Representative flow cytometry plot for gating B cell subsets from total B cells (CD19^+^CD3^−^ live cells) in the immunophenotyping experiment **D**) Proportions and absolute count of cells expressing CD69, CD95, CD11c, and CD138 within total follicular B cells (CD21^+/−^ CD23^+^) from murine splenocytes. Each point represents one mouse with bars representing the average per group. **E**) Proportions and absolute count of cells expressing CD69, CD95, CD11c, and CD138 within total marginal zone B cells (CD21^+^CD23^−^) from murine splenocytes. **F**) Proportions and absolute count of cells expressing CD69, CD95, CD11c, and CD138 within total age-associated B cells (CD21^−^CD23^−^) from murine splenocytes. **G**) Representative flow cytometry plot for B cell gating after activating splenocytes with LPS. Blue arrows mark the differentiation trajectory of these cells. **H**) Proportion of CD69^+^ cells of activated B cells (B220^+^CD138^−^) from G and MHC-II and CD69 median fluorescence intensities (MFI). **I**) Proportion of CD69^+^ cells of plasma cells (B220^−^CD138^+^) from G and CD69 MFIs. Mice with enlarged spleens were removed from spleen analyses leaving n = 8 per group. The experiment began with n = 14-15 per group. P-values were calculated using a Wilcoxon Mann-Whitney U (rank sum) test and are unadjusted for multiple comparisons.

Linear mixed effects models were used for the R-BHB and antibody titer measurements because measurements were repeated within mice, with each mouse as a random effect. For both, baseline (day 0) measurements did not differ between assigned diet groups (Wilcoxon Mann-Whitney U test, p = 0.740 for R-BHB and p = 0.880 for antibody titers), so this timepoint was excluded from modeling. For antibody titers, day 7 was also excluded because antigen-specific titers remained at background levels at this timepoint and titers are reported from the 1:1000 dilution. Values were natural log-transformed before modeling to satisfy normality and homoscedasticity, with a 0.01 mM offset for R-BHB to accommodate values at zero, and are plotted on the raw, untransformed scale. All models included timepoint as a categorical fixed effect and were fit by maximum likelihood (REML = FALSE) for likelihood ratio testing. The effect of diet was tested by comparing a model with diet and sex as fixed effects against a reduced model without diet. To test whether the effect of diet on R-BHB levels differed by sex, a model including a diet x sex interaction was compared against the corresponding additive model. P-values were calculated using likelihood ratio tests, and assumptions of the model (normality, homoscedasticity) were verified. Linear mixed effects models were generated using *lme4* version 1.1-37 in R version 4.4.2 (30).

For statistics on age-associated B cells proportions and T-bet expression after BHB treatment, linear mixed effects models were used because cells from each mouse were used at multiple BHB concentrations, with mouse as a random effect. Due to some mouse spleens having low B cell counts, not all mice are represented among all BHB concentrations (n = 4-6 mice per concentration). Models were fit to the measured values, which were natural log-transformed to satisfy normality and homoscedasticity, verified using *performance* version 0.13.0 in R version 4.4.2 (31). For visualization only, values are plotted as the change from each mouse’s untreated control (0 mM). BHB concentration was modeled as a categorical variable with the untreated condition as the reference level, and each concentration was compared to untreated control using estimated marginal means with Dunnett’s adjustment for multiple comparisons, as each concentration was compared against a common reference condition (32). Models used for these comparisons were fit by restricted maximum likelihood (REML = TRUE).

## Results

### The ketone ester diet decreases activation of B cells

To understand the effects of ketone esters on the aged immune system, aged C57BL/6JN mice (19 months old, females and males) were fed a control diet or a diet that contained the ketone ester (KE), bis-octanoyl (R)-1,3-butanediol, for 15 weeks ad libitum and spleens were collected for high-dimensional immunophenotyping (Figure 1A). This ketone ester was recently shown to be safe and well-tolerated in older adults, and to produce a sustained increase in circulating ketone body levels (26–28). We first confirmed that the ketone ester diet increased ketone body levels in the treated mice, as shown by plasma levels of the R-enantiomer of BHB (R-BHB), the primary ketone body in circulation (17). R-BHB levels were significantly higher in mice on a KE diet compared to those on a control diet (linear mixed effects model, χ^2^ (1) = 50.6, p < 0.001) (Figure 1B). There was no significant difference in the effect of the ketone ester on R-BHB levels between sexes (χ^2^ (1) = 0.067, p = 0.800) (Figure S1A). Previous studies have reported that females have higher BHB levels on a ketogenic diet compared to males, but the difference after ketone ester treatment is not significant (33–35). We also confirmed that average weights were not significantly different between mice on the control diet compared to those on the KE diet (Figure S1B).

In the spleen, B cell subsets were identified in CD19^+^CD3^−^ live cells based on expression of CD21 and CD23 (Figure 1C). There was an increase in proportion of total B cells with the ketone ester diet, but no differences in proportions nor counts of the three B cell subsets (Figure S2C-D). Bone marrow from femurs was also collected for high-dimensional immunophenotyping, but there were no changes observed in total number of cells in bone marrow, B cell progenitors, or other bone marrow-resident lineages (Figure S4B-E). This suggests that the ketone ester does not affect B cell progenitors nor B cells within the bone marrow.

Compared to those on a control diet, aged mice on the ketone ester diet showed decreased proportions of CD69^+^ and CD95^+^ populations in follicular B cells and age-associated B cells, with a trending decrease in marginal zone B cells, suggesting a decrease in activation of these subsets (Figure 1D-F). CD138 (also known as Syndecan-1) typically marks plasma cell differentiation and this marker also decreased in age-associated B cells with the ketone ester diet (Figure 1F) (36). CD138 trended toward a decrease in marginal zone B cells (Figure 1E). CD11c, which is an additional marker of age-associated B cells, was also decreased in age-associated B cells with the ketone ester diet (Figure 1F) (6,9). These results suggest that the ketone ester diet decreases activation of B cell subsets in the spleen.

Age-associated B cells have also been defined as CD11b^+/−^CD11c^+^ and by their high expression of T-bet (6,37,38). Age-associated B cells defined as CD21^−^CD23^−^ tend to be more heterogeneous as this population includes cells lacking CD11b and CD11c, whereas the majority of CD11b^+/−^CD11c^+^ B cells lack expression of CD21 and CD23 (9). To further test this decrease in activation, we also measured CD69 and CD95 expression in CD11b^+^CD11c^+^ B cells and CD11b^−^CD11c^+^ B cells as this gating also represents age-associated B cells. Within total splenic B cells, the proportions and absolute counts of CD11b^+^CD11c^+^ B cells were decreased in the ketone ester group (Figure S3A). The proportion, but not absolute counts, of CD11b^−^CD11c^+^ B cells were decreased in the ketone ester group as well. In both subsets, the proportion and absolute counts of CD95^+^ cells decreased in the ketone ester group. Only the absolute counts of CD69^+^ cells decreased, however, consistent with an overall reduction in these populations (Figure S3B-C). These results further demonstrate that the ketone ester diet decreased activation of age-associated B cells in the spleen and even decreased the proportion of age-associated B cells, as marked by expression of CD11b and CD11c.

Analysis of other splenic subsets revealed sporadic changes. There was an increase in the proportion of CD44^low^ γδ T cells in the mice with the ketone ester diet, but no change in absolute count and no changes in other T cell subsets (Table S1). While the proportion of total myeloid cells (CD11b^+^CD161^−^CD3^−^CD19^−^) was not significantly different, their absolute count decreased with the ketone ester diet. Of total myeloid cells, macrophages (F4/80^+^), monocytes (F4/80^−^Ly6C^+^Ly6G^−^), and dendritic cells (F4/80^−^Ly6C^−^Ly6G^−^ClassII^+^CD11c^high^) also significantly decreased in absolute count with the ketone ester diet, but there were no significant differences in their proportions of total myeloid cells. The total number of cells within the spleen was not significantly different between diet groups (Figure S2B). This suggests that the ketone ester diet preferentially affects B cells in the spleen as opposed to other immune cell types in aged mice. Because of the consistent effect within B cell subsets, we decided to focus on B cells.

Next, we wanted to see if the decreased activation of B cells would persist ex vivo, which would suggest cell-intrinsic effects of the ketone ester diet. To determine this, we stimulated splenocytes from the mice on the control diet or the ketone ester diet with lipopolysaccharide (LPS), a potent toll-like receptor 4 (TLR4) agonist expressed on B cells, and measured activation after four days (Figure 1A). B cells were gated based on expression of B220 and CD138. Because all cells were stimulated with LPS, they are all activated. Activated B cells (B220^+^CD138^−^) represent the earliest differentiation stage, progressing to plasmablasts (B220^+^CD138^+^) and then plasma cells (B220^−^CD138^+^) (Figure 1G).

Even after four days of LPS stimulation, we still saw a decreased proportion of the CD69^+^ population in plasma cells with a corresponding decrease in CD69 expression as shown by median fluorescence intensity (MFI) (Figure 1I). There was a trending decrease in proportion of the CD69^+^ population and CD69 MFI in activated B cells, and trending decreases in major histocompatibility complex class II (MHC-II) MFI for both subsets as well (Figure 1H-I). Because the decrease in activation persisted even after ex vivo stimulation, these results suggest that the ketone ester diet induces cell-intrinsic changes in B cells, either directly acting on them or through other immune cells.

### Antibody response after an immunization is maintained despite the decrease in B cell activation

Because we saw a decrease in B cell activation, we wanted to see if this would impair B cell function and we tested the effect of the ketone ester diet on antibody production, one of the classic functions of B cells. To analyze the effect of the ketone ester diet on antibody production during immunization, we fed aged mice with the ketone ester or control diet and immunized them with nitrophenyl-ovalbumin (NP-OVA) (Figure 2A). Plasma was collected afterward to capture the primary response to the immunization. Mice were challenged again with NP-OVA and plasma was collected to capture the memory response. In this model, the majority of the antibodies produced are specific for the hapten, NP (39,40). Total and high affinity anti-NP IgG antibodies were not different between mice on the control diet and mice on the ketone ester diet (Figure 2B). Affinity maturation, the process of improving antibody binding affinity over the course of an immune response, was also not different between groups (Figure 2C). The increase in high affinity antibodies in both diet groups also indicates properly functional germinal center reactions. These results were surprising because despite the decrease in B cell activation with the ketone ester diet, these cells were still able to produce antibodies comparably to the control diet-fed mice in response to an immunization. Taken together, these results suggest that the ketone ester diet selectively inhibits potentially deleterious, non-specific, pro-inflammatory B cells responses, as seen with decreased activation in age-associated B cells, while preserving antigen-specific B cell responses.

**Figure 2:**
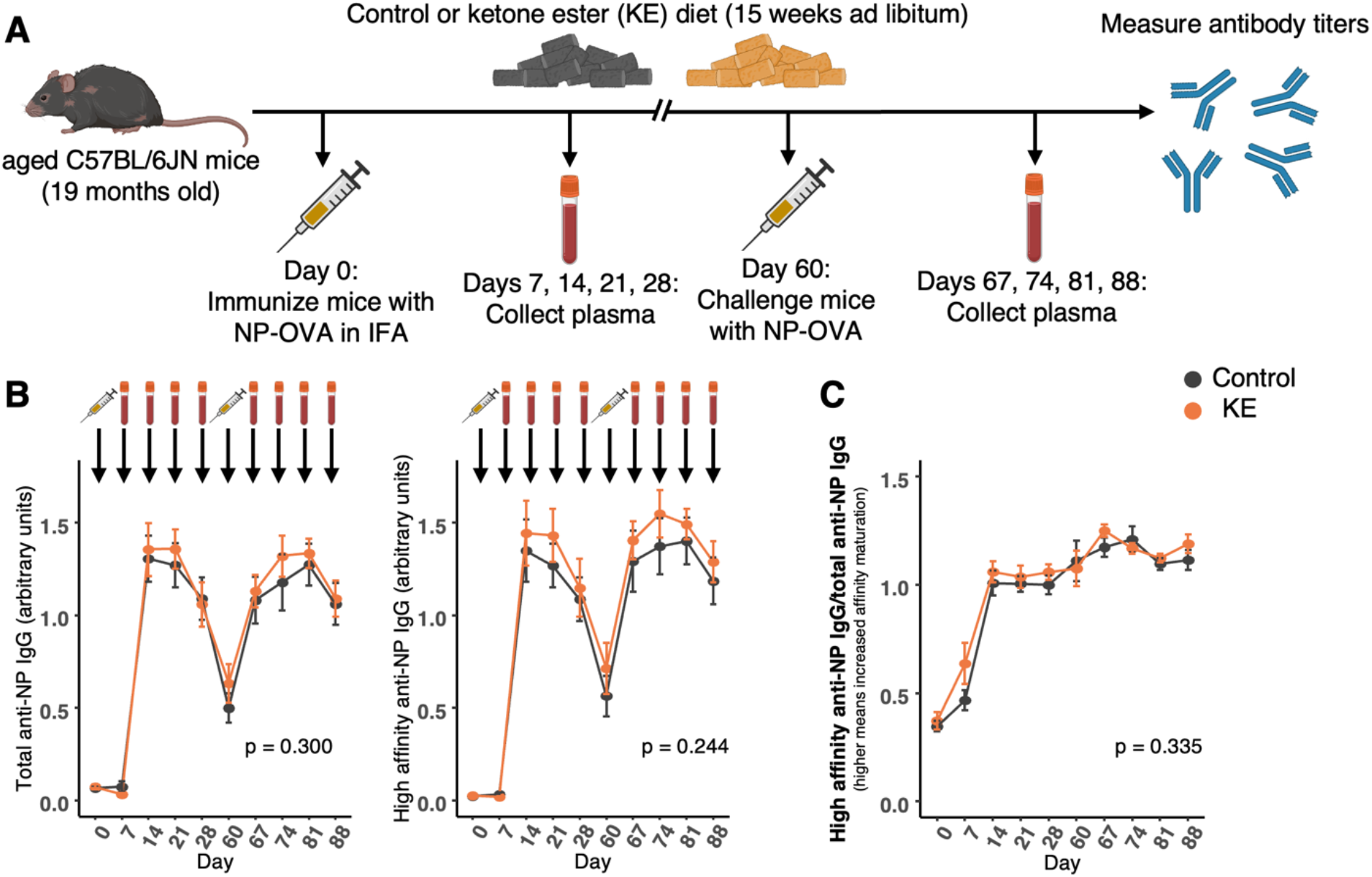
Mice on the ketone ester diet mount an immunization response despite decreased B cell activation. **A)** Experimental diagram of the immunization model. The same aged mice were immunized with nitrophenyl-ovalbumin (NP-OVA) and fed a control or ketone ester diet. Plasma was collected afterward and mice were challenged with NP-OVA before plasma was collected again. Antibody titers were measured from the plasma. **B**) Antibody titers against the hapten, NP, showing total IgG and high affinity IgG at each timepoint. Each point represents the average per group per day, and error bars represent standard error of the mean (s.e.m.). Icons above the graphs correspond to the timeline in A. **C**) Affinity maturation as measured by the ratio of high affinity anti-NP IgG antibodies to total anti-NP IgG at each timepoint. Each point represents the average per group per day, and error bars represent s.e.m. The experiment began with n = 14-15 per group. There were three non-responders per group (defined as mice that did not mount a primary response), which were excluded from the analysis. Linear mixed effects models on log-transformed data, excluding days 0 and 7 at which titers were at background levels, were used to compare mice on control diet to mice on KE diet across timepoints. Data are plotted on original scale.

### The ketone ester diet suppresses basal translation in MHC-II^low^ age-associated B cells and decreases glucose dependence

Ketone bodies are well established as an alternative energy source because they can be used for fuel in periods of low glucose through conversion into acetyl-CoA to be used in the TCA cycle. To determine how the ketone ester diet is impacting the metabolic profiles of immune cells, we performed SCENITH on the splenocytes from a subset of the same aged mouse cohort (Figure 3A). SCENITH is a method that uses puromycin incorporation as a proxy for translation and through metabolic inhibitors, can be used to determine a cell’s dependency on glucose or mitochondria for energy (41). Here, age-associated B cells were marked on their expression of CD11b and CD11c (Figure 3B).

**Figure 3:**
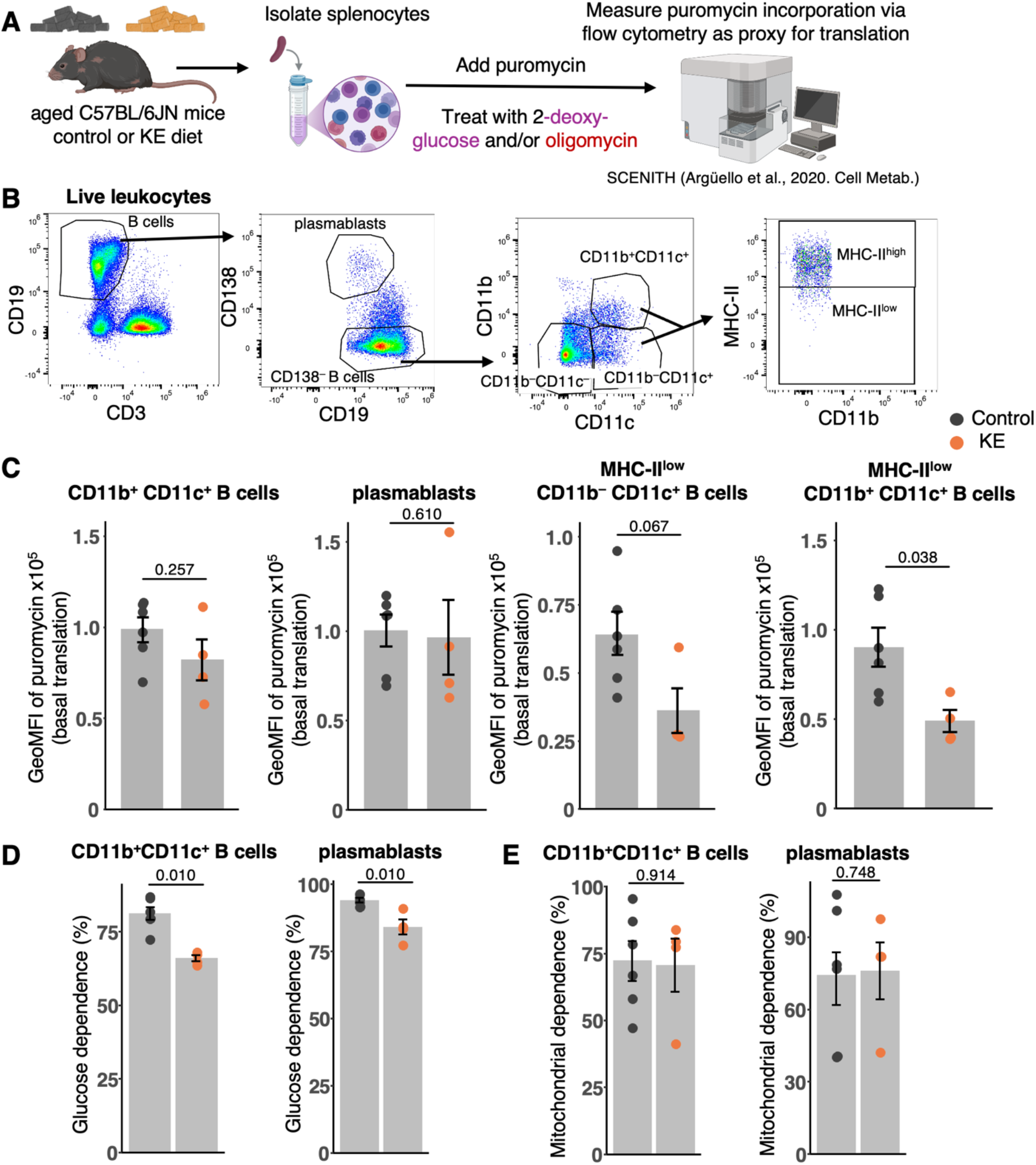
The ketone ester diet suppresses translation of MHC-IIlow age-associated B cells and reduces glucose dependence. **A**) Experimental diagram to analyze translation and metabolic usage in splenocytes from mice after control or ketone ester diet. SCENITH was used to assess puromycin incorporation as a proxy for translation (41). **B**) Gating strategy for B cells in this experiment. **C**) Geometric mean fluorescence intensities (GeoMFI) of puromycin, representing the basal translation of indicated B cell subsets. **D**) Glucose dependence of indicated B cells from mice on each diet. **E**) Mitochondrial dependence of indicated B cells from mice on each diet. SCENITH was performed on a pre-assigned subset of the aged mouse cohort (6 per group). Mice with enlarged spleens were removed from spleen analyses, leaving n = 6 for control and n = 4 for KE. Each point represents one mouse with bars representing the average per group throughout the figure. Error bars represent s.e.m. P-values were calculated using a Wilcoxon Mann-Whitney U (rank sum) test and are unadjusted for multiple comparisons.

First, we determined basal translation by measuring the level of puromycin incorporation. Compared to those from control, splenocytes from the mice on the ketone ester diet showed no difference in translation in bulk CD11b^+^CD11c^+^ B cells or in plasmablasts (Figure 3C). When we further gated these cells based on MHC-II expression, however, the ketone ester diet decreased translation in MHC-II^low^ CD11b^+^CD11c^+^ B cells, with a trending decrease in MHC-II^low^ CD11b^−^CD11c^+^ B cells (Figure 3C). This suggests that the ketone ester inhibits translation within the MHC-II^low^ compartment of age-associated B cells rather than across the bulk population.

Compared to control, the ketone ester diet decreased glucose dependence in CD11b^+^CD11c^+^ cells and plasmablasts (Figure 3D). This is consistent with current knowledge of how ketone bodies can promote oxidative phosphorylation. However, mitochondrial dependence was not different in these cells, suggesting that the ketone ester selectively inhibits glucose usage without impacting mitochondria (Figure 3E). These data suggest that the ketone ester diet inhibits translation of age-associated B cells expressing low levels of MHC-II while also reducing glucose dependence, though these measurements were made in fewer mice than the immunophenotyping analyses.

### The ketone body β-hydroxybutyrate is sufficient to suppress B cell translation

We were curious to see if the suppression of B cell translation from the ketone ester diet was due to the effect of ketone bodies themselves. To test this, we isolated splenocytes from aged wildtype mice (24-month-old males), stimulated them overnight with LPS, and treated with β-hydroxybutyrate (BHB) before running SCENITH, gating B cell subsets based on CD21 and CD23 expression (Figure 4A-B).

**Figure 4:**
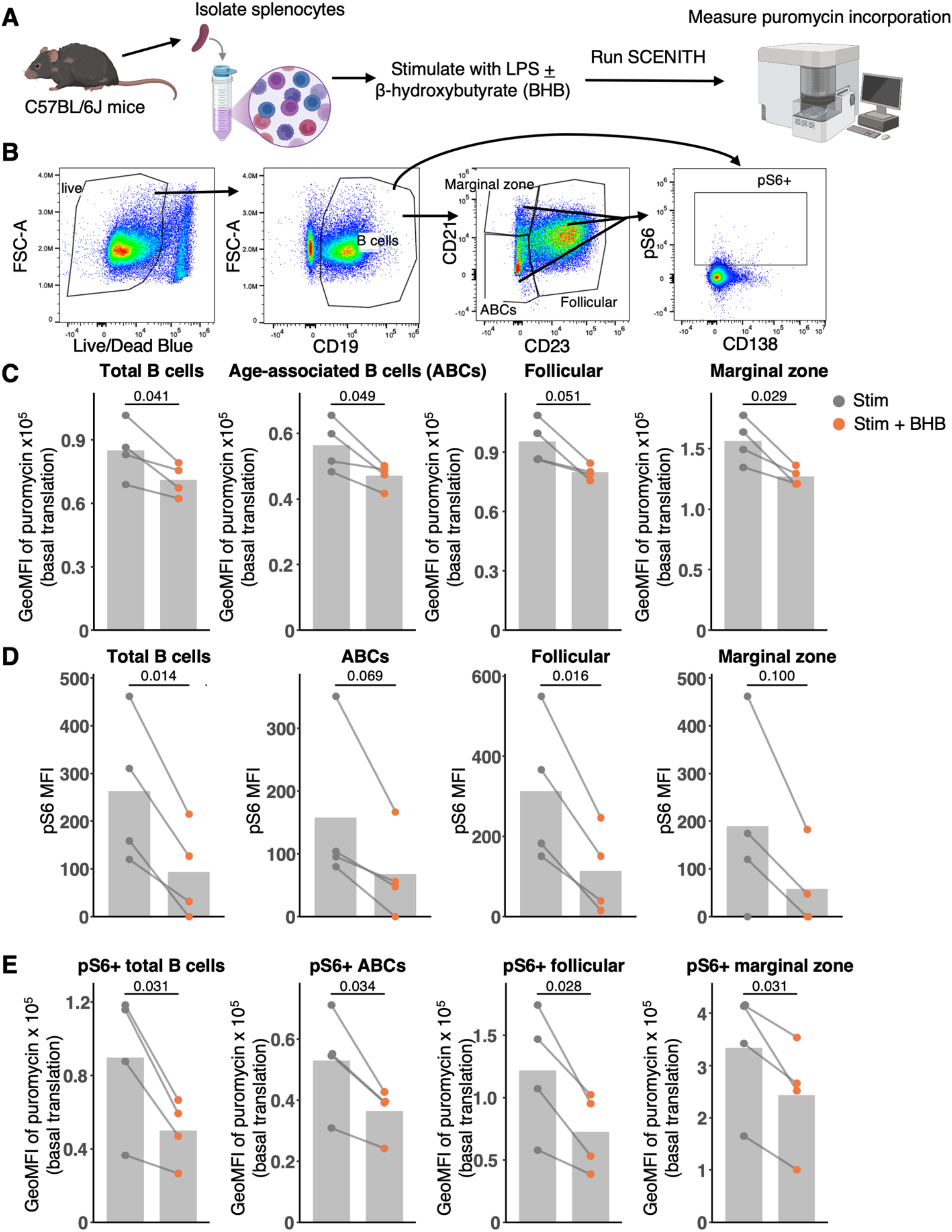
The ketone body β-hydroxybutyrate is sufficient to suppress B cell translation. **A)** Experimental diagram to analyze translation and metabolic usages of splenocytes after treatment with β-hydroxybutyrate (BHB) with LPS stimulation using SCENITH. **B**) Gating strategy for B cells in this experiment. **C)** Geometric mean fluorescence (GeoMFI) intensities of puromycin, representing the basal translation of indicated B cell subsets after LPS stimulation with and without BHB treatment. Each point represents one mouse with bars representing the average per condition and lines represent cells from the same mouse (n = 4). **D**) Median fluorescence intensities (MFIs) of pS6 on B cell subsets. **E**) GeoMFIs of puromycin for phosphorylated-S6 (pS6)^+^ populations of B cells after LPS stimulation with and without BHB treatment. P-values in C were calculated from one-sided paired T-tests as the a priori hypothesis based on Figure 3 was that BHB would decrease translation. P-values in D and E were calculated from two-sided paired T-tests as the pS6 analyses were exploratory and the direction of any effect was not predicted.

Stimulation with BHB compared to stimulation alone decreased translation in virtually all B cell subsets: total B cells, age-associated B cells, marginal zone B cells, and follicular B cells (trending decrease) (Figure 4C). These results demonstrate that BHB alone is sufficient to suppress B cell translation. BHB treatment did not alter glucose or mitochondrial dependence (Figure S5A-B), further suggesting that neither the ketone ester nor BHB increases mitochondrial dependence in B cells.

The mTOR pathway is classically known for protein synthesis, cell growth, and proliferation. The ribosomal protein S6 is downstream of mTORC1, and phosphorylation of S6 (pS6) by mTORC1-dependent S6 kinase enhances translation (42,43). Because of these decreases in translation, we were curious if the mTOR pathway was involved. To determine if mTOR signaling was altered, we measured phosphorylation of S6 in this experiment. Expression of pS6, measured by median fluorescence intensity (MFI), was decreased in total B cells and follicular B cells with BHB treatment, while it was trending toward a decrease in age-associated B cells and marginal zone B cells (Figure 4D). The proportion of pS6^+^ cells significantly decreased in marginal zone B cells with BHB treatment but not in the other B cell subsets (Figure S5C). BHB treatment also inhibited translation in all pS6^+^ B cell subsets, indicating that BHB still inhibits translation through other mechanisms apart from the mTOR-S6K axis (Figure 4E). These results suggest that BHB inhibits translation in B cells both through inhibition of mTOR and through other downstream or parallel mechanisms.

### β-hydroxybutyrate reduces T-bet expression in age-associated B cells

T-bet is the signature transcription factor in age-associated B cells that drives their autoimmune functions (7,44). In T cells, T-bet and mTOR are connected as mTORC1 was shown to promote phosphorylation of T-bet (45). Because of this connection between mTOR and T-bet and the importance of T-bet in age-associated B cells, we were curious if BHB impacts T-bet expression. We isolated B cells from spleens of aged wildtype mice (24-month-old males) and treated them with a cocktail to differentiate them into age-associated B cells: CL097 (a TLR7/8 agonist), IL-21, IFNγ, IgM, and CD40 (46). We also treated with varying concentrations of β-hydroxybutyrate (BHB) and incubated cells for 3 days before measuring age-associated B cell proportion and T-bet expression (Figure 5A, Figure S6). Here, we looked at three types of gating for age-associated B cells: CD21^−^CD23^−^ B cells, CD11b^−^CD11c^+^ B cells, and CD11b^+^CD11c^+^ B cells.

**Figure 5:**
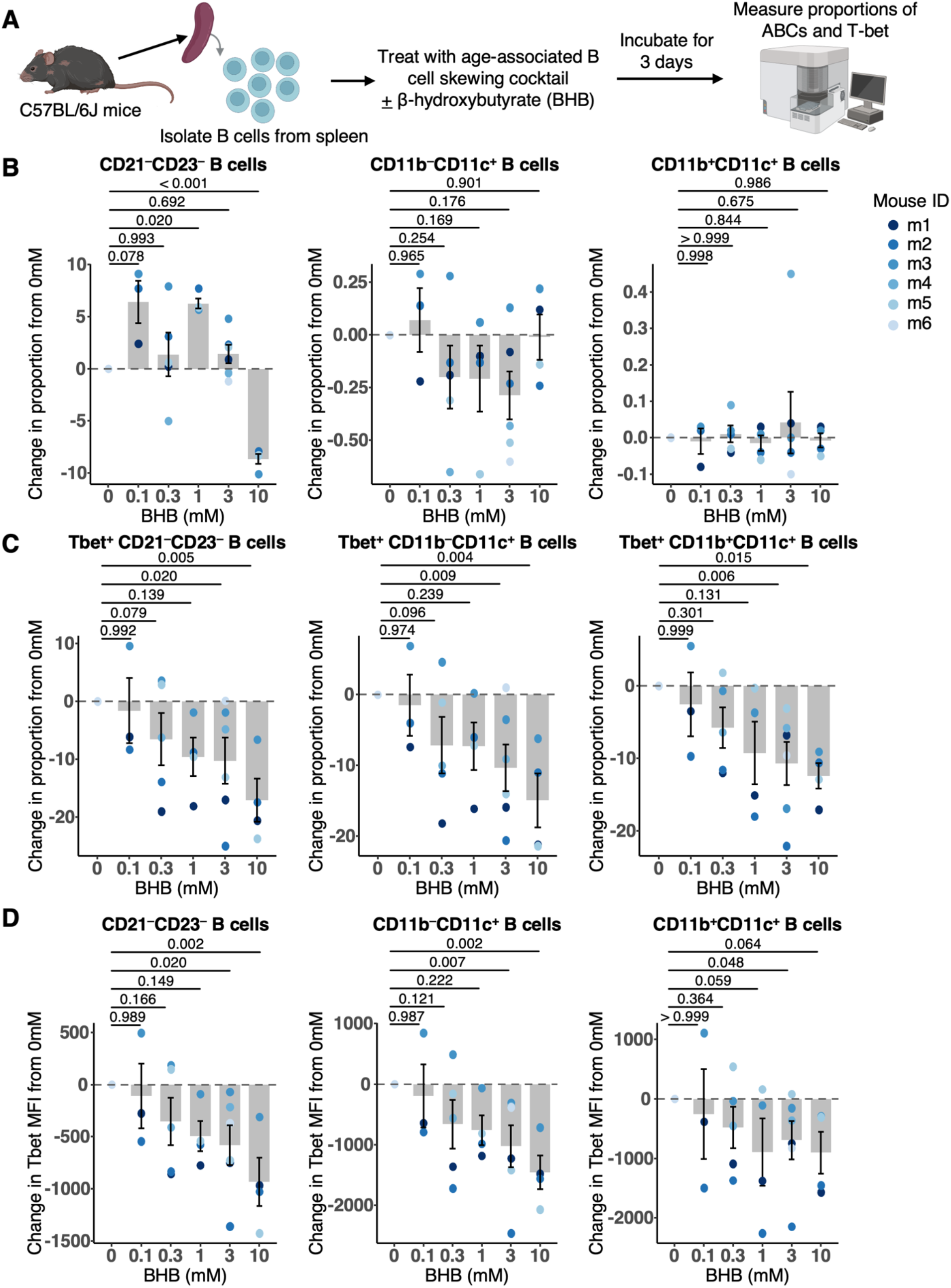
β-hydroxybutyrate decreases T-bet expression in age-associated B cells (ABCs) **A)** Experimental diagram for age-associated B cell differentiation with β-hydroxybutyrate (BHB) treatment (0 – 10 mM). **B**) Change in proportions of CD21^−^CD23^−^ B cells, CD11b^−^CD11c^+^ B cells, CD11b^+^CD11c^+^ B cells relative to each mouse’s untreated control (0 mM). **C**) Change in proportions of T-bet^+^ populations within each subset relative to each mouse’s untreated control (0 mM). **D**) Change in median fluorescence intensity (MFI) of T-bet within each subset relative to each mouse’s untreated control (0 mM). Each point represents one mouse at that condition and points are colored per mouse (n = 4-6 mice per dose). Bar graphs represent the average per condition. Error bars represent s.e.m. P-values were calculated from linear mixed effects models fit to natural log-transformed raw values with mouse as a random effect, comparing each BHB concentration to untreated control with a Dunnett’s post-hoc test. Plotted values show the change from each mouse’s untreated control and were not used for modeling.

Compared to untreated control, BHB significantly decreased the proportion of CD21^−^CD23^−^ B cells at a concentration of 10 mM (Figure 5B). However, it significantly increased their proportion at 1 mM and trended toward an increase at 0.1 mM. As mentioned before, CD21^−^CD23^−^ age-associated B cells are heterogeneous, while most CD11b^+/−^CD11c^+^ age-associated B cells lack expression of CD21 and CD23 (9). At all concentrations, BHB did not change the proportion of CD11b^−^CD11c^+^ B cells nor CD11b^+^CD11c^+^ B cells. Because CD11b^+/−^CD11c^+^ expression identifies age-associated B cells more specifically than CD21^−^CD23^−^ B cells, which captures a more heterogeneous population, the absence of any change from BHB in the CD11c^+^ subsets indicates that BHB does not impact the generation of ABCs. In all three age-associated B cell subsets, BHB significantly decreased the proportion of T-bet^+^ cells at the higher concentrations of 3 and 10 mM (Figure 5C). The proportion of T-bet^+^ cells trended toward a decrease at 0.3 mM BHB in CD21^−^CD23^−^ B cells and CD11b^−^CD11c^+^ B cells. T-bet expression measured by MFI was also significantly decreased with BHB treatment at concentrations of 3 and 10 mM in CD21^−^ CD23^−^ B cells and CD11b^−^CD11c^+^ B cells (Figure 5D). It was significantly decreased at a concentration of 3 mM in CD11b^+^CD11c^+^ B cells and trended toward a decrease at 1 and 10 mM (Figure 5D). These results show that BHB inhibits the expression of T-bet in age-associated B cells at high concentrations. Whether this occurs at the circulating BHB levels achieved by the ketone ester diet remains to be determined.

In summary, the ketone ester diet in aged mice decreased expression of activation markers in splenic B cells, especially in age-associated B cells, without impairing antigen-specific antibody production following immunization. Glucose dependence was also decreased in age-associated B cells, measured in a smaller subset of the cohort. BHB suppressed B cell translation by inhibiting the mTORC1 pathway as well as through other mechanisms beyond S6 phosphorylation. BHB also inhibited the expression of T-bet in age-associated B cells at high concentrations, raising the possibility that this contributes to the suppressed activation seen with the ketone ester diet.

## Discussion

In this study, we found that the ketone ester diet in aged mice inhibited B cell activation, especially in age-associated B cells. Despite this decrease in activation, B cells were still able to produce antibodies in response to NP-OVA immunization, showing that this reduced activation did not impair antibody production. This suggests that the ketone ester selectively inhibits the baseline activation of B cells, while preserving the ability of these cells to mount a functional antigen-specific response when challenged, such as during immunization. This distinction matters for aging, where the goal is to dampen chronic, antigen-independent activation that accompanies inflammaging without blunting the immune responses that older individuals rely on. We also provide evidence that BHB is sufficient to decrease B cell translation, likely through inhibition of the mTOR pathway. Translation was also inhibited in B cells with activated mTOR despite seeing a decrease in mTOR activity with BHB, suggesting that there are other unknown mechanisms through which BHB inhibits translation. Lastly, BHB reduced the expression of T-bet in age-associated B cells in vitro. Because T-bet drives the activation and effector functions of these cells, this reduction suggests a potential mechanism by which the ketone ester diet may dampen age-associated B cell activation in vivo, though this was observed at BHB concentrations above those measured in circulation. These results highlight novel effects of ketone esters and ketone bodies on B cells, especially in the context of aging. BHB is known to have anti-inflammatory properties, especially in innate immune cells, such as inhibiting the NLRP3 inflammasome (18). Our work further supports the anti-inflammatory effects of ketone bodies by adding a new mechanism within B cells.

Age-associated B cells are implicated in aging and autoimmune diseases by secreting proinflammatory cytokines and autoimmune antibodies (6,7,9). The connection between fasting and ketogenic diets and age-associated B cells is unclear, but fasting-mimicking diets and ketogenic diets have been reported to improve autoimmune diseases in humans and in mouse models (14–16,47). Interestingly, mTOR has also been implicated in autoimmune diseases. T-bet^+^ CD11c^+^ memory B cells (thought to be a human equivalent of age-associated B cells) are expanded in systemic lupus erythematosus (SLE) and have overactive mTORC1 compared to classical memory B cells (48). mTOR inhibitors are currently in clinical trials to test whether they can improve autoimmune disease, and sirolimus has shown promising results (49). Ketone esters may be able to recapitulate some of these improvements in autoimmunity and offer a more accessible treatment because they can work as a supplement instead of requiring an entire diet change. Whether ketone esters can ameliorate autoimmunity in autoimmune mouse models or in clinical trials remains an important question for future investigation.

Apart from fueling the TCA cycle, BHB also has other epigenetic and signaling functions. It is a histone deacetylase (HDAC) inhibitor and a post-translational modification itself (known as β-hydroxybutyrylation) on lysine residues on histones, non-histone proteins, and even antibodies (21,50–52).

Our data showed that BHB was sufficient to suppress translation of B cells, likely mediated through inhibition of mTOR signaling. mTORC1 has been shown to be important for B-cell responses, specifically for class switch recombination and germinal center formation (53). Ketone body metabolism and mTORC1 signaling are interconnected. mTORC1 activity suppresses ketogenesis and BHB has been shown to inhibit mTORC1 (with the exception of muscle) (54). A recent report showed that β-hydroxybutyrylation on lysine 349 of RagC inhibits the interaction of mTOR with RagC, thus suppressing the pathway (55). BHB also activates AMPK, which also inhibits mTORC1 (56,57). This knowledge is consistent with the decrease in pS6 expression seen in our data. Because mTORC1 has been shown to promote T-bet expression in T cells (45), the reduction in T-bet that we observed with BHB may also occur downstream of mTOR inhibition, though this remains to be tested. BHB has been reported to affect other immune cells through these non-metabolic functions, so it would be interesting to investigate whether BHB acts on age-associated B cells through these other mechanisms (21,58,59).

One important consideration is the relationship between the BHB concentrations used in vitro and those achieved in vivo. Plasma R-BHB in mice on the ketone ester diet reached 0.5 to 1 mM, whereas the effects on T-bet expression were observed at 3 and 10 mM. Several factors make it difficult to directly compare the two. Plasma samples were collected during the early light phase, several hours after the dark-phase feeding period, and may reflect steady-state rather than peak concentrations in ad libitum-fed animals. Exposure to the ketone ester also differed from BHB treatment: mice received the ketone ester for 15 weeks, while cells were treated for three days, comparing chronic to acute treatments. The concentrations required to reduce T-bet in vitro exceed those we measured in circulation, and whether this mechanism operates at physiological ketone body levels remains to be determined.

A recent paper reported that fasting and a ketogenic diet through BHB actually deplete long-lived plasma cells (LLPCs; defined by high expression of CD138 and TACI/CD267) from the bone marrow, leading to impaired immune memory (58). In contrast, we found no differences in proportions of CD138^+^TACI^+^ cells in the bone marrow of mice fed the ketone ester diet (Figure S4C). Zhu et al. also reported reduced antigen-specific antibody titers after immunization, whereas our cohort showed no difference in antibody response after NP-OVA immunization (Figure 2B-2C). These conflicting results may arise from multiple methodological differences between the studies. First, the two studies analyzed different phases of the humoral response. Zhu et al. waited 8 weeks after immunization to measure antibodies, allowing enough time for long-lived plasma cells to develop and home to the bone marrow. They then measured antibody titers for another 16 weeks and largely saw differences in the later timepoints. In our study, we measured antigen-specific antibodies up to about 12 weeks after immunization. Thus, the two studies are analyzing different time periods: the shorter term response to immunization compared to the longer term one. It is possible that our null result reflects the timing of when we measured rather than an absence of an effect on the long-lived plasma cells. Secondly, the ketogenic interventions differed. We fed mice a ketone ester diet, while their study used fasting (alternate day and prolonged), a ketogenic diet, and intraperitoneal injection of BHB. The prolonged fasting regimen lasted up to 3 days, which is rather intense for mice considering they can lose about 20% of their body weight after 2 days of fasting (60). Recall that the ketone ester diet is meant to supplement ketone bodies and recapitulates this one aspect of ketogenic diets and fasting. It is unclear how BHB exposure from the ketone ester diet compares to that achieved by injection in their study. Lastly, we examined aged mice whereas they studied young mice (8-12 weeks old), potentially showing age-dependent effects of ketone bodies on B cells. Future studies should focus on understanding the whole longitudinal view of ketone bodies’ effects on the development of immune memory and how timing, dosage, and age play key roles.

This study has several limitations. Antibody titers did not differ between diet groups, but the precision of this comparison is limited by cohort size (95% confidence intervals on the diet effect: fold changes of 0.88 to 1.51 for total anti-NP IgG and 0.88 to 1.65 for high affinity anti-NP IgG). The SCENITH experiment was performed on a subset of the mouse cohort (6 control and 4 ketone ester), and the resulting metabolic findings should be considered exploratory pending confirmation in a larger cohort. Mice with splenomegaly were excluded from spleen analyses, reducing the analyzed cohort. More ketone ester than control mice were excluded (5 versus 2). This difference was not statistically significant (Fisher’s exact test, p = 0.405), but the small numbers give the test little power to detect an imbalance. Thus, we cannot exclude the possibility that this differential exclusion of ketone ester mice contributed to the lower activation observed in that group. Immunophenotyping comparisons were unadjusted for multiple comparisons and exploratory. While individual markers may not survive correction, the decrease in activation was consistent in direction across CD69, CD95, CD138, and CD11c, and across follicular, marginal zone, and age-associated B cell subsets. It is unclear if the amount of food consumed differed between groups, and time since last feeding was not recorded for individual mice. Because circulating BHB varies with feeding, this contributes variability to individual measurements. All mice were sampled non-fasted within the same window of the light phase, however, and weights were not different between groups (Figure S1B), making large differences in energy intake unlikely. The diets differed not only by inclusion of the ketone ester, but by carbohydrate and fat percentage: the control diet was 77.7% carbohydrate and 12.6% fat, while the KE diet was 51.1% carbohydrate and 39.2% fat. The ketone ester is a high fat supplement, and we provide evidence that BHB alone is sufficient to suppress B cell translation and T-bet expression in vitro (Figures 4-5), supporting a ketone body-mediated mechanism. A high-fat control diet would be needed to formally separate the contribution of dietary fat from that of the ketone ester. Another limitation is that we were not able to compare immunophenotyping by sex, as some mice died before the final timepoint and enlarged spleens were removed from analyses. Given that autoimmune diseases are more prevalent in females than males (4), future work should determine whether the ketone ester diet affects immune cell subsets differently by sex.

## Conclusions

In conclusion, the ketone ester, bis-octanoyl (R)-1,3-butanediol, decreased activation of B cells in aged mice, particularly age-associated B cells, without impairing the antibody response to immunization. The primary ketone body BHB was sufficient to suppress translation in B cells, and to reduce T-bet expression in age-associated B cells at high concentrations. Our study highlights a new immunoregulatory mechanism of ketone bodies in B cells and positions ketone esters as a promising intervention worth testing in models for age-related autoimmunity.

## List of Abbreviations

KE: ketone ester
BHB: β-hydroxybutyrate
R-BHB: R-β-hydroxybutyrate
ABC: age-associated B cell
MFI: median fluorescence intensity
GeoMFI: geometric mean fluorescence intensity
NP-OVA: nitrophenyl-ovalbumin
LPS: lipopolysaccharide
IFA: incomplete Freund’s adjuvant
MHC-II: major histocompatibility complex II
SCENITH: single-cell energetic metabolism by profiling translation inhibition

## Declarations

## Ethics approval and consent to participate

All mice were maintained in accordance with the guidelines set forth by the National Institutes of Health, and all experimental protocols were approved by the Buck Institute Institutional Animal Care and Use Committee (IACUC). The Buck Institute is accredited by the Association for Assessment and Accreditation of Laboratory Animal Care (AAALAC).

## Consent for publication

Not applicable

## Availability of data and materials

All data generated or analyzed during this study are included in this published article and its supplementary information files (SourceData.xlsx).

## Competing interests

E.V. and J.C.N. are co-founders, hold stock, and are co-inventors on patents licensed to Component Health, Selah Therapeutics, and BOPZ. J.C.N. is on the scientific advisory board on Junevity, Inc. E.V. is the founder of Napa Therapeutics.

## Funding

This work was supported by funding from the following sources: National Institute on Aging & National Institutes of Health R01AG068025 (E.V., J.C.N.), U01AI180158 (E.V.), R01AG067333 (J.C.N.), T32AG052374 (A.A.-F.) and T32AG000266 (M.N.); Longevity Impetus (J.C.N.); Hevolution Foundation HF-PART-23-1422047 (E.V., J.C.N.); the Michael Antonov Foundation (E.V.); and Buck Institute institutional funds (E.V., J.C.N.).

## Authors’ contributions

A.A.-F., R.T., H.G.K., M.N., and J.C.N. designed the experiments. A.A.-F. and R.T. performed the experiments with the help of M.N., R.R.R., R.K., D.S., and M.M.K. A.A.-F., R.T., and H.G.K. analyzed the experiments. A.A.-F. drafted and wrote the manuscript. All authors reviewed and edited the manuscript. J.C.N. and E.V. supervised the study.

## Acknowledgments

Experimental diagrams were made with the help of BioRender.com

## Additional files

**Additional file 1.** SupplementaryFiguresandTables_ImmunityandAgeing.docx – Supplementary Figures 1-6 and Supplementary Tables 1-3

**Additional file 2.** SourceData.xlsx – Source data for all figure panels.

## Notes

### Summary of Updates

Revised throughout, including new title, a restructured abstract with keywords, and expanded background and discussion sections. New figures (Figure 5 and Supplementary Figure 3) added. Methods, Results, and Discussion sections updated.

