## Supplementary Figures and Tables for "Ketone ester supplementation reduces age-associated B cell activation in aged mice without impairing antibody responses"

**Supplementary Figures 1-6**

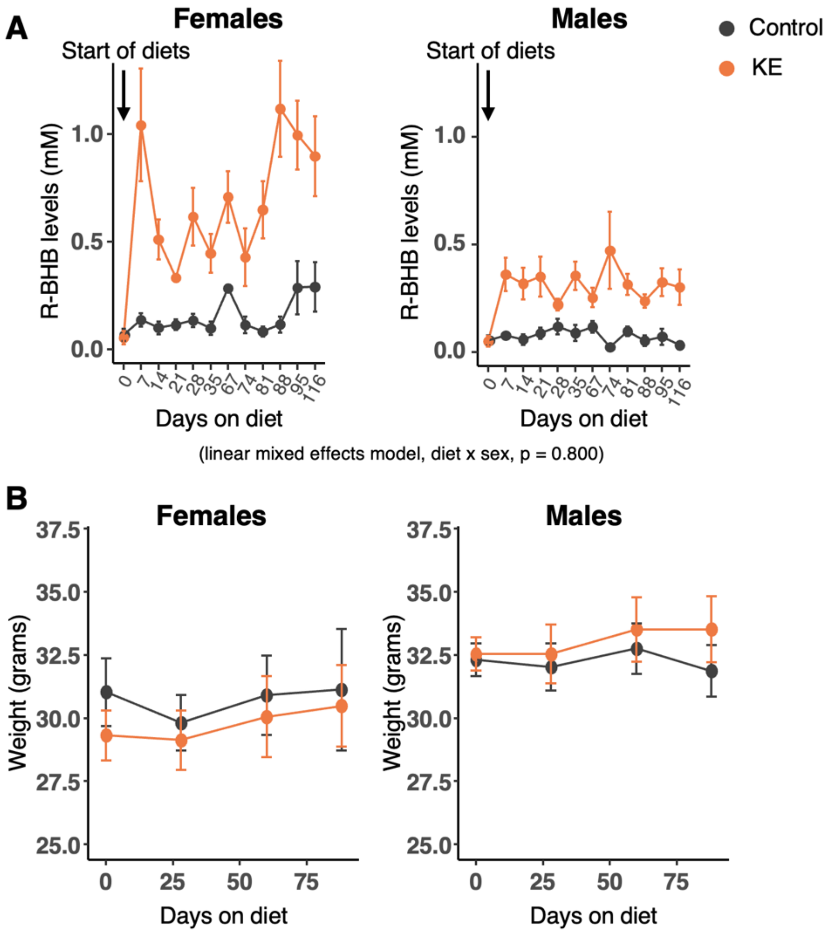


**Supplementary Figure 1: Validation of the ketone ester diet**

**A)** R-BHB levels of mice throughout the experiment separated by sex (mean ± s.e.m.). A linear mixed effects model on log-transformed R-BHB levels, excluding baseline (day 0), was used to compare females on KE to males on KE. Data are plotted on original scale. **B**) Weights of mice throughout the duration of the experiment on the diets separated by sex. Weights were measured throughout the diet but simplified to representative points here. KE stands for ketone ester. The experiment began with n = 14-15 per diet group and ended with n = 10-13 per diet group due to mice that did not survive.

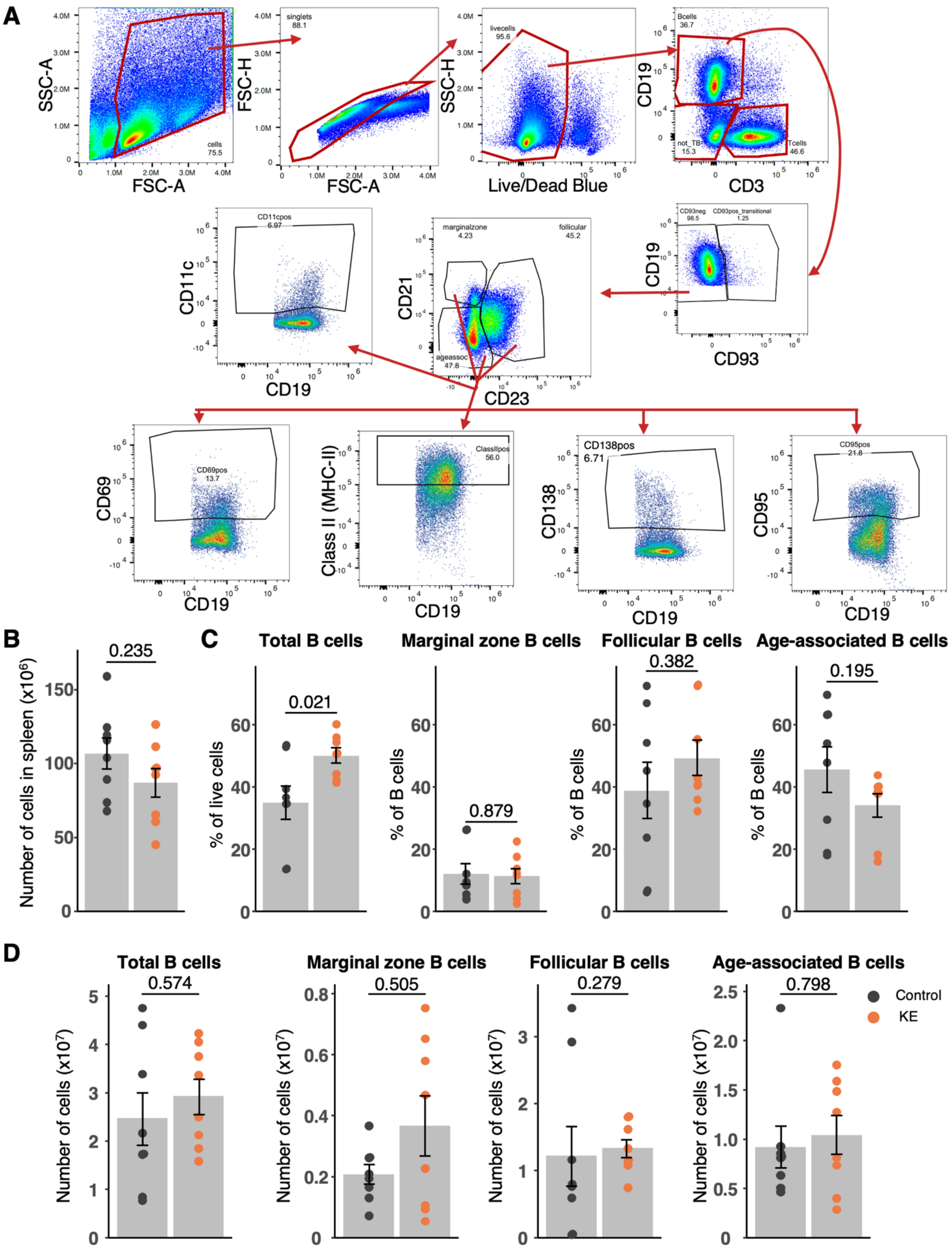

A
SSC-A
FSC-A
cells 75.5
FSC-H
singlets 88.1
SSC-H
livecells 95.6
Live/Dead Blue
CD19
Bcells 36.7
not_TB 15.3
Tcells 46.6
CD3
CD93neg 98.5
CD93pos_transitional 1.25
CD93
CD21
marginalzone 4.23
follicular 45.2
ageassoc 47.8
CD23
CD11c
CD11cpos 6.97
CD69
CD69pos 13.7
Class II (MHC-II)
ClassIIpos 56.0
CD138
CD138pos 6.71
CD95
CD95pos 21.8
B
Number of cells in spleen (x10⁶)
0.235
C
Total B cells
0.021
% of live cells
Marginal zone B cells
0.879
% of B cells
Follicular B cells
0.382
Age-associated B cells
0.195
D
Total B cells
0.574
Number of cells (x10⁷)
Marginal zone B cells
0.505
Follicular B cells
0.279
Age-associated B cells
0.798
Control
KE

**Supplementary Figure 2: Total amounts of splenic B cell subsets were not different with the ketone ester diet**

**A)** Gating strategy for B cell immunophenotyping in the spleen. **B**) Total number of cells in the spleen. **C**) Proportions of total B cells as a percentage of total live cells in the spleen. Proportions of B cell subsets (marginal zone, follicular, and age-associated B cells) as percentages of total B cells in the spleen. **D**) Absolute cell count of total B cells and B cell subsets in the spleen. KE stands for ketone ester. Each point represents one mouse with bars representing the average per group throughout the figure. Error bars represent s.e.m. Mice with enlarged spleens were removed from the analysis leaving n = 8 per group. P-values were calculated using a Wilcoxon Mann-Whitney U (rank sum) test and are unadjusted for multiple comparisons.

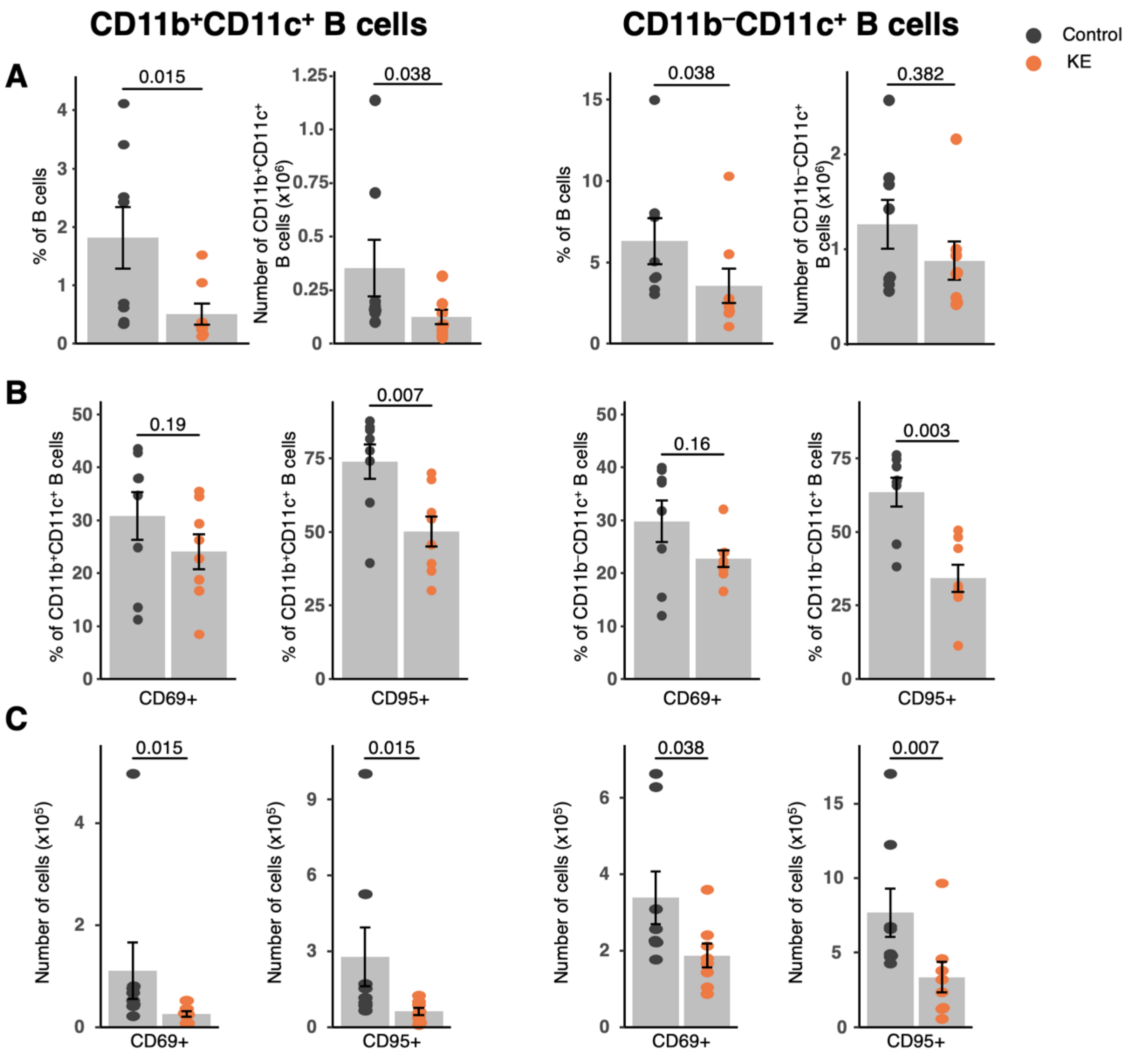


**Supplementary Figure 3: The ketone ester diet decreases CD11b/CD11c B cell subsets in the spleen and their activation.**

**A)** Proportions within total B cells and absolute counts of CD11b$^{+}$CD11c$^{+}$ B cells and CD11b$^{-}$CD11c$^{+}$ B cells from murine splenocytes. **B**) Proportion of cells expressing CD69 and CD95 within CD11b$^{+}$CD11c$^{+}$ B cells and CD11b$^{-}$CD11c$^{+}$ B cells. **C**) Absolute counts of cells expressing CD69 and CD95 within CD11b$^{+}$CD11c$^{+}$ B cells and CD11b$^{-}$CD11c$^{+}$ B cells. Each point represents one mouse. Bar graphs represent the average per group and error bars represent s.e.m. Mice with enlarged spleens were removed from spleen analyses leaving n = 8 per group. The experiment began with n = 14-15 per group. P-values were calculated using a Wilcoxon Mann-Whitney U (rank sum) test and are unadjusted for multiple comparisons.

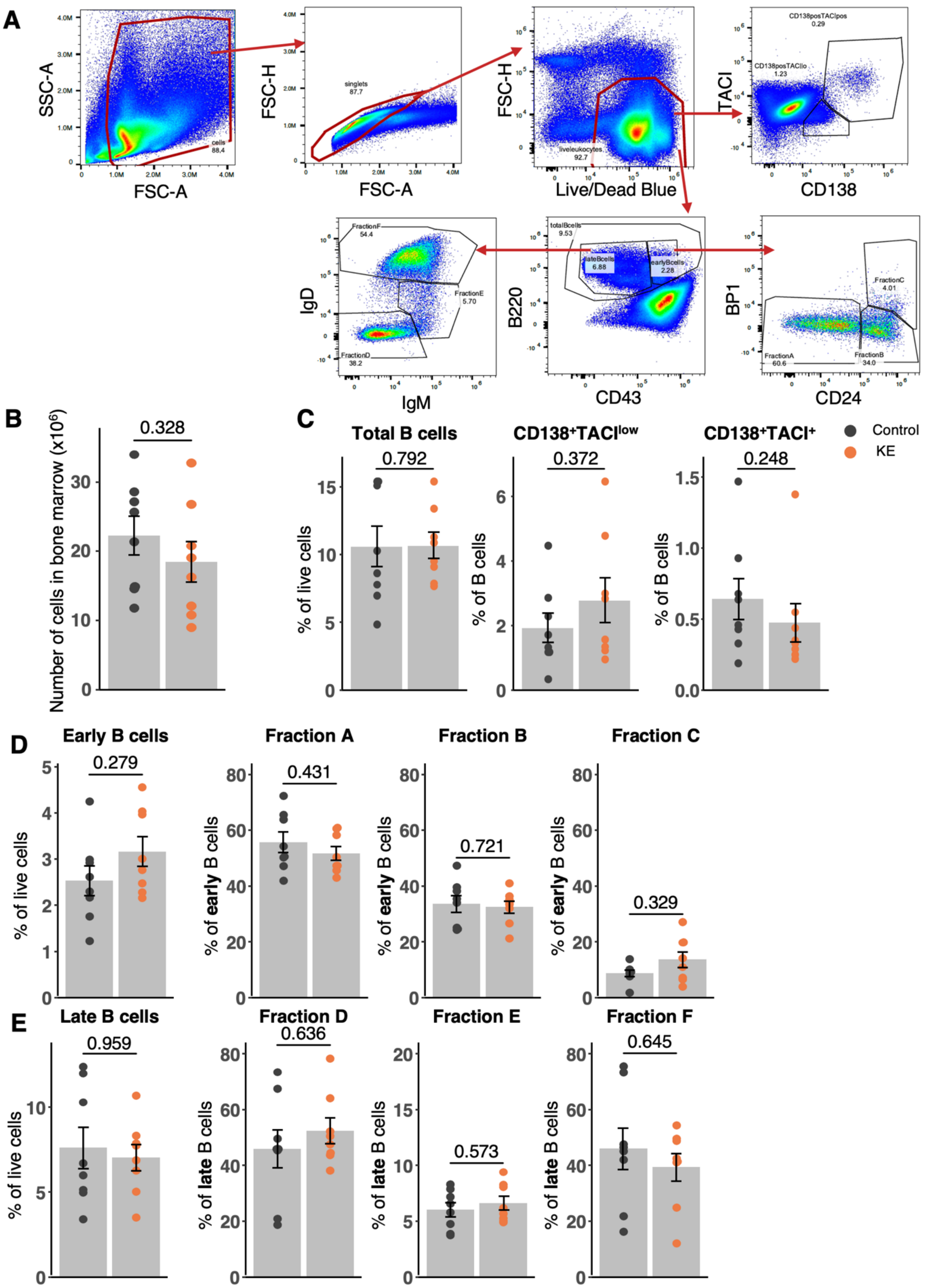
A
SSC-A
FSC-A
cells
88.4
FSC-H
singlets
87.7
Live/Dead Blue
liveleukocytes
92.7
TACI
CD138
CD138posTACIpos
0.29
CD138posTACIlo
1.23
IgD
IgM
FractionF
54.4
FractionE
5.70
FractionD
38.2
B220
CD43
totalBcells
9.53
lateBcells
6.88
earlyBcells
2.28
BP1
CD24
FractionC
4.01
FractionA
60.6
FractionB
34.0
B
Number of cells in bone marrow (x10^6)
0.328
C
Total B cells
0.792
% of live cells
CD138+TACIlow
0.372
% of B cells
CD138+TACI+
0.248
% of B cells
Control
KE
D
Early B cells
0.279
% of live cells
Fraction A
0.431
% of early B cells
Fraction B
0.721
% of early B cells
Fraction C
0.329
% of early B cells
E
Late B cells
0.959
% of live cells
Fraction D
0.636
% of late B cells
Fraction E
0.573
% of late B cells
Fraction F
0.645
% of late B cells

**Supplementary Figure 4: The ketone ester diet did not change B cell subsets in the bone marrow**

**A)** Gating strategy for B cell immunophenotyping in the bone marrow. **B**) Total number of cells in the bone marrow. **C**) Proportions of total B cells as a percentage of total live cells in the bone marrow. Proportions of relevant CD138 and TACI/CD267 subsets as a percentage of total B cells in the bone marrow. **D**) Proportion of early B cells as a percentage of total live cells. Proportions of the three fractions as percentages of early B cells. **E**) Proportion of late B cells as a percentage of total live cells. Proportions of the three fractions as percentages of late B cells. Each point represents one mouse with bars representing the average per group throughout the figure. Error bars represent s.e.m. n = 8 per group. P-values were calculated using a Wilcoxon Mann-Whitney U (rank sum) test and are unadjusted for multiple comparisons.

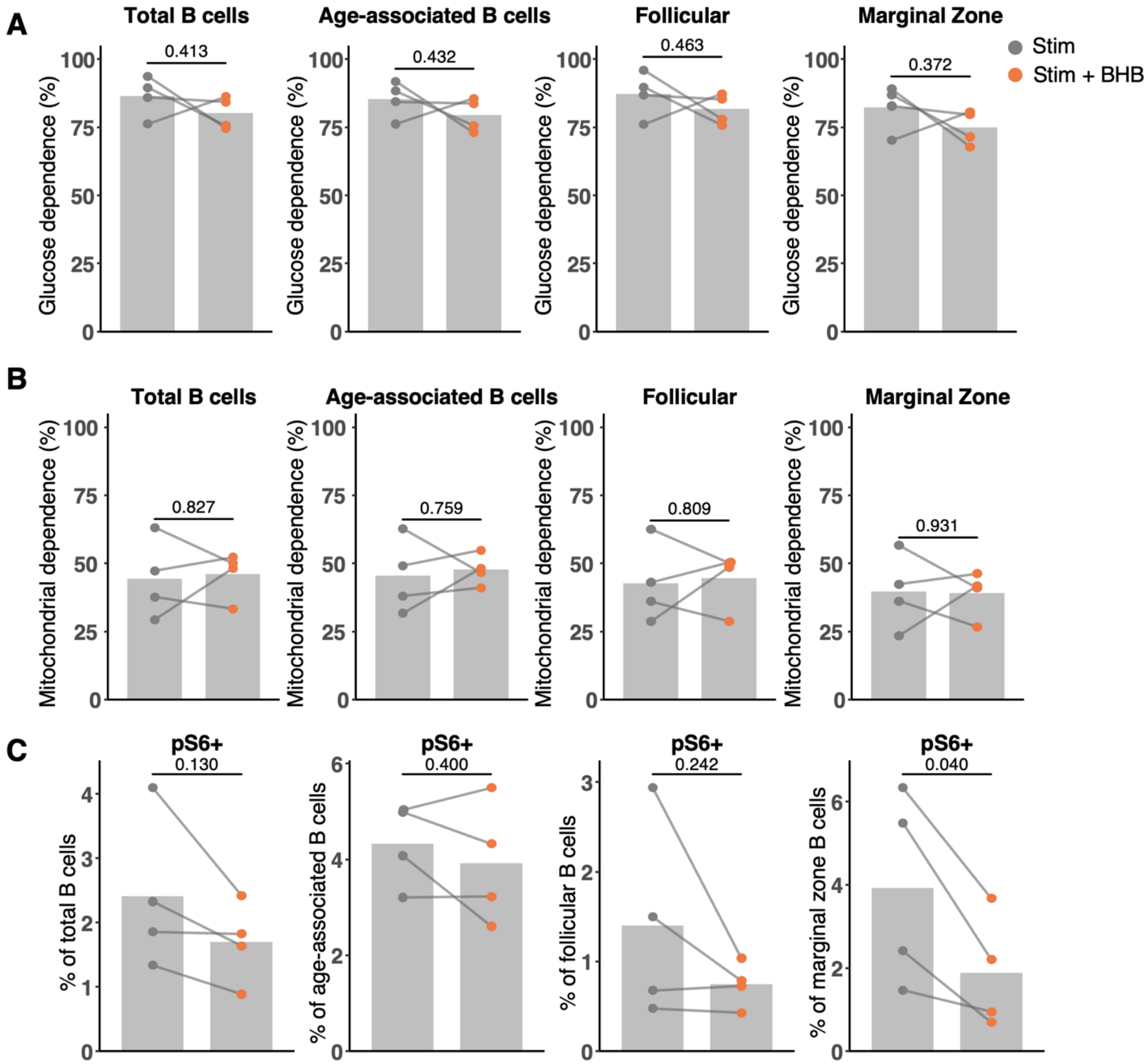


**Supplementary Figure 5: BHB does not alter glucose dependence, mitochondrial dependence, or pS6-positive proportions in B cells**

**A)** Glucose dependence of indicated B cells after LPS stimulation with and without BHB treatment. **B**) Mitochondrial dependence of indicated B cells after LPS stimulation with and without BHB treatment. **C**) Proportion of pS6+ populations within total B cells, age-associated B cells, follicular B cells, and marginal zone B cells. Each point represents one mouse with bars representing the average per condition and lines represent cells from the same mouse (n = 4). P-values were calculated from two-sided paired T-tests as the pS6 analyses were exploratory and the direction of any effect was not predicted.

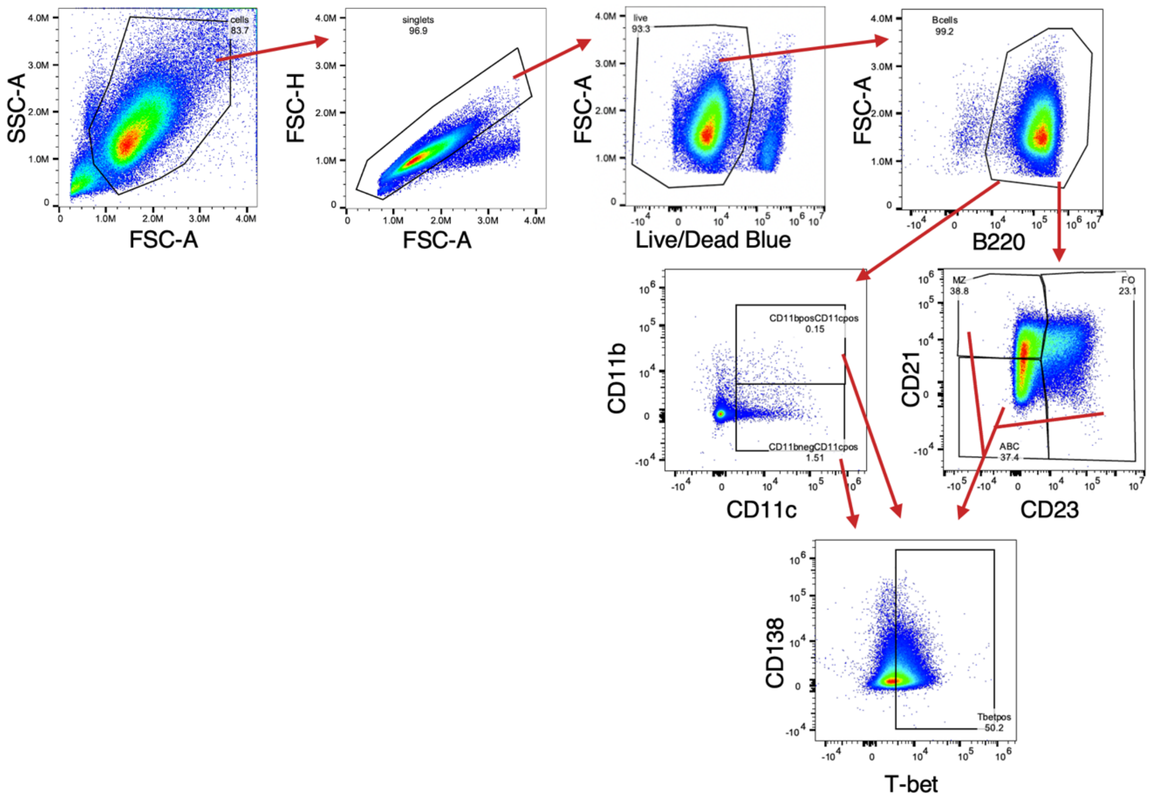


**Supplementary Figure 6: Gating strategy for Figure 5**

Gating strategy for age-associated B cell subsets shown in Figure 5.

**Supplementary Tables 1-3**

**Supplementary Table 1: Additional immunophenotyping changes in the spleen with the ketone ester diet**

P-values were calculated using a Wilcoxon Mann-Whitney U (rank sum) test and are unadjusted for multiple comparisons. Fold changes represent the average ratio between the ketone ester (KE) diet and control diet.

| **Subset** | **Proportion** | | **Absolute cell count** | |
|---|---|---|---|---|
| | **p-value** | **Fold change (KE/Control)** | **p-value** | **Fold change (KE/Control)** |
| $CD44^{low}$ γδ T cells | 0.028 | 1.53 | > 0.999 | 2.49 |
| total myeloid cells | 0.398 | 0.90 | 0.050 | 0.596 |
| monocytes | 0.083 | 0.50 | 0.038 | 0.14 |
| macrophages | 0.074 | 1.31 | 0.050 | 0.596 |
| dendritic cells | 0.380 | 1.17 | 0.050 | 0.57 |

**Supplementary Table 2: Flow cytometry panel for spleen immunophenotyping**

| Target | Fluorochrome | Vendor | Clone | Catalog Number |
|---|---|---|---|---|
| CD3 | APC-Fire 810 | BioLegend | 17A2 | 100267 |
| CD4 | BV650 | BioLegend | RM4-5 | 100545 |
| CD8 | PerCP | BioLegend | 53-6.7 | 100732 |
| CD11b | PerCP-Cy5.5 | BioLegend | M1/70 | 101227 |
| CD11c | PE-Fire 810 | BioLegend | QA18A72 | 161106 |
| CD19 | PE-Dazzle 594 | BioLegend | 6D5 | 115553 |
| CD21/35 | BV421 | BioLegend | 7E9 | 123422 |
| CD23 | Alexa Fluor 700 | BioLegend | B3B4 | 101631 |
| CD25 | Spark NIR 685 | BioLegend | PC61 | 102069 |
| CD44 | BV570 | BioLegend | IM7 | 103037 |
| CD45 | BV510 | BioLegend | 30-F11 | 103137 |
| CD62L | BV711 | BioLegend | MEL-14 | 104445 |
| CD69 | PE-Cy7 | BioLegend | H1.2F3 | 104512 |
| CD93 | FITC | BioLegend | AA4.1 | 136507 |
| CD95 | Alexa Fluor 647 | BioLegend | SA367H8 | 152619 |
| CD127 | PE-Fire 700 | BioLegend | S18006K | 158215 |
| CD138 | BV785 | BioLegend | 281-2 | 142534 |
| CXCR5 | BUV395 | BD Biosciences | 2G8 | 563960 |
| F4/80 | BV480 | BD Biosciences | T45-2342 | 565635 |
| I-A/I-E (MHC-II) | BUV496 | BD Biosciences | 2G9 | 750171 |
| ICOS | Pacific Blue | BioLegend | C398.4A | 313521 |
| IgM | APC-Cy7 | BioLegend | RMM-1 | 406515 |
| KLRG1 | PE | BioLegend | 2F1/KLRG1 | 138407 |
| Ly-6C | BUV805 | BD Biosciences | RB6-8C5 | 741920 |
| Ly-6G | BUV563 | BD Biosciences | 1A8 | 612921 |
| NK1.1 (CD161) | BV605 | BioLegend | PK136 | 108739 |
| PD-1 | BUV615 | BD Biosciences | RMP1-30 | 752354 |
| TCRγδ | APC | BioLegend | GL3 | 118115 |
| Viability | LIVE DEAD Blue | Thermo Fisher | - | L34962 |

**Supplementary Table 3: Flow cytometry panel for bone marrow immunophenotyping**

| Target | Fluorochrome | Vendor | Clone | Catalog Number |
|---|---|---|---|---|
| CD3 | APC-Fire 810 | BioLegend | 17A2 | 100267 |
| CD11b | PerCP-Cy5.5 | BioLegend | M1/70 | 101227 |
| CD16/32 | BUV395 | BD Biosciences | 190909 | 747953 |
| CD19 | BV570 | BioLegend | 6D5 | 115535 |
| CD24 | FITC | BioLegend | QA20A91 | 114305 |
| CD34 | BV421 | BioLegend | MEC14.7 | 119321 |
| CD43 | PE-Cy7 | BioLegend | S11 | 143209 |
| CD45 | BUV563 | BD Biosciences | 30-F11 | 612924 |
| CD48 | BV711 | BioLegend | HM48-1 | 103439 |
| CD117 | BV650 | BioLegend | 2B8 | 105853 |
| CD127 | PE | BioLegend | S18006K | 158204 |
| CD135 | PE-Cy5 | BioLegend | A2F10 | 135312 |
| CD138 | BV785 | BioLegend | 281-2 | 142534 |
| CD150 | PE-Dazzle 594 | BioLegend | TC15-12F12.2 | 115935 |
| B220 | PE-Fire 810 | BioLegend | RA3-6B2 | 103287 |
| BP-1 | BV510 | BD Biosciences | BP-1 | 745061 |
| F4/80 | BV480 | BD Biosciences | T45-2342 | 565635 |
| IgD | Alexa Fluor 647 | BioLegend | 11-26c.2a | 405707 |
| IgM | APC-Cy7 | BioLegend | RMM-1 | 406515 |
| Ly-6C | BUV805 | BD Biosciences | RB6-8C5 | 741920 |
| Ly-6G | PerCP | BioLegend | 1A8 | 127653 |
| NK1.1 (CD161) | BV605 | BioLegend | PK136 | 108739 |
| Sca-1 | Pacific Blue | BioLegend | D7 | 108119 |
| TACI (CD267) | APC | eBioscience | ebio8F10-3 | 17-5942-82 |
| Ter-119 | Alexa Fluor 700 | BioLegend | TER-119 | 116220 |
| Viability | LIVE DEAD Blue | Thermo Fisher | - | L34962 |
